# Innate immunity pathways activate cell proliferation after penetrating traumatic brain injury in adult *Drosophila*

**DOI:** 10.1101/2021.09.01.458615

**Authors:** Khailee Marischuk, Kassi L. Crocker, Shawn Ahern-Djamali, Grace Boekhoff-Falk

## Abstract

We are utilizing an adult penetrating traumatic brain injury (PTBI) model in Drosophila to investigate regenerative mechanisms after damage to the central brain. We focused on cell proliferation as an early event in the regenerative process. To identify candidate pathways that may trigger cell proliferation following PTBI, we utilized RNA-Seq. We find that transcript levels for components of both Toll and Immune Deficiency (Imd) innate immunity pathways are rapidly and highly upregulated post-PTBI. We then tested mutants for the NF-κB transcription factors of the Toll and Imd pathways, Dorsal-related immunity factor (Dif) and Relish (Rel) respectively. We find that loss of either or both Dif and Rel results in loss of cell proliferation after injury. We then tested canonical downstream targets of *Drosophila* innate immune signaling, the antimicrobial peptides (AMPs), and find that they are not required for cell proliferation following PTBI. This suggests that there are alternative targets of Toll and Imd signaling that trigger cell division after injury. Furthermore, we find that while AMP levels are substantially elevated after PTBI, their levels revert to near baseline within 24 hours. Finally, we identify tissue-specific requirements for Dif and Rel. Taken together, these results indicate that the innate immunity pathways play an integral role in the regenerative response. Innate immunity previously has been implicated as both a potentiator and an inhibitor of regeneration. Our work suggests that modulation of innate immunity may be essential to prevent adverse outcomes. Thus, this work is likely to inform future experiments to dissect regenerative mechanisms in higher organisms.

## Introduction

The scarcity of neural stem cells in the adult brain represents an apparent barrier to therapeutic neural regeneration in humans. Treatments for neurodegenerative diseases and neural injuries thus have focused primarily on the transplantation of stem cells (Gurgo et al., 2002; Vishwakarma et al., 2014). However, transplants are complex, costly, inefficient, and often have undesirable side effects, including tumor formation. For these reasons, therapies that activate endogenous regenerative mechanisms would be preferable. Indeed, regeneration from resident cells is a growing area of research, but remains technically challenging. We are utilizing a *Drosophila* model to investigate brain regeneration. Adult *Drosophila* brains, like mammalian ones, have few neural stem cells. However, *Drosophila* are amenable to genetic techniques not easily utilized in higher organisms.

Using a penetrating traumatic brain injury (PTBI) model, we find that cells in the adult *Drosophila* brain are stimulated to proliferate upon injury (Crocker et al., 2021). The dividing cells give rise to both new glia and new neurons near the injury site. However, the mechanism that activates proliferation post-PTBI is not known. Preliminary studies indicated that the innate immune signaling pathways might be involved. *Drosophila* have an innate immune system, but no adaptive immune system (De Gregorio et al., 2002). The *Drosophila* innate immune system is composed of the Toll and Immune deficiency (Imd) pathways (Leclerc and Reichhart, 2004). The Toll pathway is stimulated by microbial peptidoglycans or damage-associated molecular patterns (DAMPs) via the activation of an extracellular ligand, Spätzle (Spz) (Leclerc and Reichhart, 2004). A signaling cascade leading to cleavage of Spz is activated by the Peptidoglycan recognition protein-SA (PGRP-SA)/Gram-negative bacteria binding protein 1 (GNBP1) complex or the Gram-negative bacteria binding protein 3 (GNBP3) and involves multiple proteases, particularly Spätzle-Processing Enzyme (SPE), to activate immune signaling. Cleaved Spz is recognized by the Toll receptors at the cell membrane, triggering a pathway that results in the degradation of Cactus, a negative regulator of two nuclear factor-kappa B (NF-κB) transcription factors, Dorsal and Dorsal-related immunity factor (Dif). Both Dorsal and Dif regulate immune signaling in embryos and larvae, while Dif appears to be more important in adults (Rutschmann et al., 2000). Translocation of Dif to the nucleus results in the transcriptional upregulation of canonical immune signaling target genes, which encode antimicrobial peptides (AMPs) (Lemaitre et al., 1997). AMPs can kill microbes, thus removing pathogenic threats. The Imd pathway also is activated by microbial peptidoglycans, but the relevant receptors are either freely distributed in the hemolymph, such as PGRP-LE, or located in the cellular membrane, such as PGRP-LC (Myllymaki et al., 2014). Binding of microbial peptidoglycans causes a complex of Imd, Death related ced-3 (Dredd), and Fas-associated death domain (Fadd) to be recruited. This complex activates downstream kinases, leading to the phosphorylation and cleavage of a third NF-κB transcription factor, Relish (Rel). Once Rel has been activated, it migrates to the nucleus to induce the expression of AMPs and other immune-related genes (Myllymaki et al., 2014).

It is thought that immune signaling in the brain is detrimental to the tissue, because overexpression of AMPs in glia results in neurodegeneration, and Rel activity underlies the neurodegeneration in a fly model of ataxia-telangiectasia (Cao et al., 2013; Petersen et al., 2012). In the context of brain injury, penetrating injury to the *Drosophila* optic lobe is sufficient to activate expression of AMPs and other genes associated with the immune response (Sanuki et al., 2019). Closed head traumatic brain injury (TBI) also activates innate immune signaling, leading to neurodegeneration (Sanuki et al., 2019). Previous studies thus implicate Toll and Imd activation in cell and/or organismal death following brain injury.

In contrast, innate immune signaling in embryos and larvae can stimulate wound closure and promote cell survival (Capilla et al., 2017; Parisi et al., 2014). Linking beneficial roles of innate immune signaling to adult neural regeneration has not been done previously in *Drosophila*. Based on initial studies correlating upregulation of innate immunity with cell proliferation following penetrating brain injury, we hypothesized that the Toll and Imd pathways promote neurogenesis after brain injury. Consistent with this hypothesis, immune activation and inflammation can promote either neurogenesis or neurodegeneration in mammals, depending on the microenvironment around the injury and the relative scale of the inflammatory response (Kyritsis et al., 2014).

Our work confirms previous *Drosophila* brain injury studies in demonstrating that immune signaling pathways are upregulated after penetrating brain injury (Sanuki et al., 2019). However, in addition, we find that the activation of Toll and Imd signaling through the NF-κB transcription factors, Dif and Rel, is required to trigger cell proliferation following a PTBI. Further we find that this signaling is required in specific cell types both within and surrounding the brain. Nonetheless, the canonical targets of innate immune signaling, the AMPs, are not required for cell proliferation post-PTBI. We conclude that there are other targets of Toll and Imd signaling that are essential for this particular injury response. Based on the transience of AMP upregulation, we propose that rapid modulation of the immune response may be essential to forestall secondary injury and neurodegeneration. Taken together, this work illustrates that one of major regulators of proliferation after injury is the innate immune system and that the precise spatial and temporal control of immunity cascades may trigger the initial steps required for neural regeneration after injury.

## Materials & Methods

### Fly Stocks and Rearing

All flies were reared at 25°C on a standard cornmeal-sugar medium. The following stocks were obtained from the Bloomington Drosophila Stock Center: #458 (*p{GawB}elav[C155]*), #25374 (*y[1] w[*]; P{Act5C-GAL4-w}E1/CyO*), #7415 (*w*^*1118*^; *P{w[+m*]=GAL4}repo/TM3, Sb*^*1*^), #9458 (*w*^*1118*^; *Rel*^*E38*^ *e*^*s*^), #28943 (*y*^*1*^ *v*^*1*^; *P{y[+t7*.*7] v[+t1*.*8]=TRiP*.*HM05154}attP2*), #30141 (*w*^*1118*^; *P{w[+mC]=Hml-GAL4*.*Delta}3*), #30513 (*y*^*1*^ *sc*^*\**^ *v*^*1*^ *sev*^*21*^; *P{y[+t7*.*7] v[+t1*.*8]=TRiP*.*HM05257}attP2*), #34559 (*P{ry[+t7*.*2]=Dipt2*.*2-lacZ}1, P{w[+mC]=Drs-GFP*.*JM804}1, y*^*1*^ *w*^*\**^; *Dif*^*1*^ *cn*^*1*^ *bw*^*1*^), #36558 (*P{ry[+t7*.*2]=Dipt2*.*2-lacZ}1, P{w[+mC]=Drs-GFP*.*JM804}1, y*^*1*^ *w*^*\**^; *Dif*^*2*^ *cn*^*1*^ *bw*^*1*^), #51635 (*y*^*1*^ *w*^*\**^; *P{w[+m*]=nSyb-GAL4S}3*), #55707 (*P{ry[+t7*.*2]=Dipt2*.*2-lacZ}1, P{w[+mC]=Drs-GFP*.*JM804}1, y*^*1*^ *w*^*\**^), #55714 (*Rel*^*E20*^), and #58814 (*y*^*1*^ *w*^*\**^; *P{w[+mC]=yolk-GAL4}2*). Double mutant *Dif*^*1*^*/Dif*^*2*^; *Rel*^*E20*^*/Rel*^*E38*^ stocks were generated in our laboratory using the stocks listed above. The AMP deletion stocks were generously provided by the Lemaitre lab: Group A deletion strain (lacking *Defensin*), Group B deletion strain (lacking *Drosocin, Attacin A, Attacin B, Attacin D, Diptericin A*, and *Diptericin B*), Group C deletion strain (lacking *Metchnikowin* and *Drosomycin*), and Group ABC deletion strain. Full descriptions of these stocks are available in Hanson *et al*., 2019 (Hanson et al., 2019).

### Penetrating Traumatic Brain Injury

To induce PTBI, we used thin metal needles (∼12.5 μm diameter tip, 100 μm rod; Fine Science Tools) sterilized in 70% ethanol to penetrate the head capsule of CO_2_-anesthetized adult flies. Injured flies were transferred back to our standard sugar food for recovery and aging. The same injury method was applied to flies for 5-ethynyl-2’-deoxyuridine (EdU) labeling, except flies were fed 50 mM EdU in 10% sucrose solution on a size 3 Whatman filter for six hours before injury, then allowed to recover on the same solution for 24 hours. For immunohistochemistry experiments, each fly was unilaterally injured in the right hemisphere of the central brain. For molecular experiments, each fly was injured bilaterally.

### RNA-Seq

RNA-Seq samples were isolated from heads of males four hours after PTBI to both hemispheres of the central brain and from the heads of age- and sex-matched controls without injury. RNA-seq workflow integrated service was provided by ProteinCT Biotechnologies LLC (Madison, WI). Libraries were prepared using the TruSeq strand-specific mRNA sample preparation system (Illumina). The final library was generated by further purification and amplification with PCR, and quality checked using a Bioanalyzer 2100. The libraries were then sequenced (single end 50 bp reads) using the Illumina HiSeq2500, with six samples (three control and three injured) per lane, for a total of over twenty million reads per sample. Next, the fast QC program was used to verify raw data quality of the Illumina reads. The *Drosophila melanogaster* genome and gene annotations were downloaded from FlyBase and used for mapping. The raw sequence reads were mapped to the genome using Subjunc aligner from Subread, with the majority of the reads (over 98% for all samples) aligned to the genome. The alignment bam files were compared against the gene annotation GFF file, and raw counts for each gene were normalized using the voom method from the R Limma package. Once normalized, they were then used for differential expression analysis. Hierarchical clustering was used to indicate sample and gene relationships. In the overall heatmap, each column is a sample, and each row represents the scaled expression values for one gene (blue is low, red is high). In this clustering there were 367 genes with differential expression of 2-fold or greater between the two groups. 259 of those were upregulated, and 108 were downregulated with a false discovery rate (FDR) <0.05.

### Immunohistochemistry

Brains were dissected in PBS and fixed in a 3.7% formaldehyde in a PEM (100 mM piperazine-N,N’-bis(2-ethanesulfonic acid) [PIPES], 1 mM EGTA, 1 mM MgSO_4_) solution for 20 minutes at 25°C. Fixed brain samples were washed in PT (PBS [phosphate-buffered saline] and 1% Triton X-100), blocked with 2% BSA in PT solution (PBT), and then incubated with primary antibodies overnight at 4°C in PBT. Following primary incubation, the samples were washed with PT and then incubated overnight in secondary antibody at 4°C. The next day, samples were washed in PT, stained with DAPI (1:10,000, ThermoFisher) for 8 minutes, and mounted in Vectashield anti-fade mountant (Vector Labs). The primary antibodies used in this study are rabbit anti-PH3 (1:500, Santa Cruz Biotechnology, Inc). Secondary antibodies used are anti-rabbit 568 (1:400, ThermoFisher). For EdU labeling, brains were dissected, processed, and antibody stained as described above using buffers without azide prior to the Click-IT® reaction. EdU detection was performed by following the manufacturer’s protocol (InVitrogen). All slides were imaged using a Nikon A1RS system and analyzed using the Nikon NIS Elements software. Cell counting was done both manually and using the Nikon NIS-Elements software to detect regions of interest (ROIs) with a threshold of over 1000 and an area of at least 10μm.

### Quantitative Real-Time PCR

Transcript levels of target genes were measured by quantitative real-time PCR (qRT-PCR) using method described in Ihry et al. 2012 (Ihry et al., 2012). RNA was isolated from appropriately-staged animals using the RNeasy Plus Mini Kit (Qiagen). cDNA was synthesized from 40 to 400 ng of total RNA using the SuperScript III First-Strand Synthesis System (Invitrogen). qPCR was performed on a Roche 480 LightCycler using the LightCycler 480 DNA SYBR Green I Master kit (Roche). In all cases, samples were run simultaneously with three independent biological replicates for each target gene, and *Rp49* was used as the reference gene. To calculate changes in relative expression, the Relative Expression Software Tool was utilized. The following primers were used for qRT-PCR: *DptA* Forward: 5’-GCTGCGCAATCGCTTCTAC-3’ & Reverse: 5’-TGGTGGAGTGGGCTTCATG-3’ (Chakrabarti et al., 2014); *Drs* Forward: 5’-CTGGGACAACGAGACCTGTC-3’ & Reverse: 5’-ATCCTTCGCACCAGCACTTC-3’ (Fly Primer Bank); *CecB* Forward: 5’-TTGTGGCACTCATCCTGG-3’ & Reverse: 5’-TCCGAGGACCTGGATTGA-3’ (Kleino et al., 2008); *Dro* Forward: 5’-GCACAATGAAGTTCACCATCGT-3’ & Reverse: 5’-CCACACCCATGGCAAAAAC-3’ (Tsai et al., 2008); *Rp49* Forward: 5’-CCAGTCGGATCGATATGCTAA-3’ & Reverse: 5’-ACGTTGTGCACCAGGAACTT-3’ (Denton et al., 2009).

## Results

### Innate Immunity Genes Are Upregulated After PTBI

As reported in Crocker *et al*. (Crocker et al., 2021), we have developed a novel method to study neural regeneration. We use a fine sterile needle to inflict penetrating traumatic brain injury (PTBI) to the mushroom body (MB) region of the young adult brain (Crocker et al., 2021). After injury, we see a regenerative process that begins with an increase in the number of proliferating cells (Crocker et al., 2021). These newly created cells then differentiate into neurons and glia that can then integrate into the brain to functionally repair the damage (Crocker et al., 2021). In order to understand how regeneration occurs, we chose to focus on one of the first steps in the process: cell proliferation (**Fig. 1A**). To identify genes that may be involved in activating proliferative mechanisms in the brain, we analyzed changes in gene expression after PTBI via RNA-Sequencing (RNA-Seq). We used RNA isolated from the heads of males four hours after double injury to both mushroom bodies and from age-matched controls of the same sex. There were 367 genes with differential expression of 2-fold or greater between the injured and control groups. Of these, 259 were upregulated, and the remaining 106 were downregulated, with a false discovery rate (FDR) of <0.05. Gene Ontology (GO) terms for the upregulated genes in our injured samples were identified to determine what biological processes were affected upon injury. The single most highly enriched set of genes were involved defense and immune response, with 27% of identified GO Terms falling into this category (**Fig. 1B**). Components of both major innate immune pathways were upregulated four hours after injury along with the canonical downstream readouts of immune activation, the AMPs (**Fig. 1B**).

**Figure 1.**
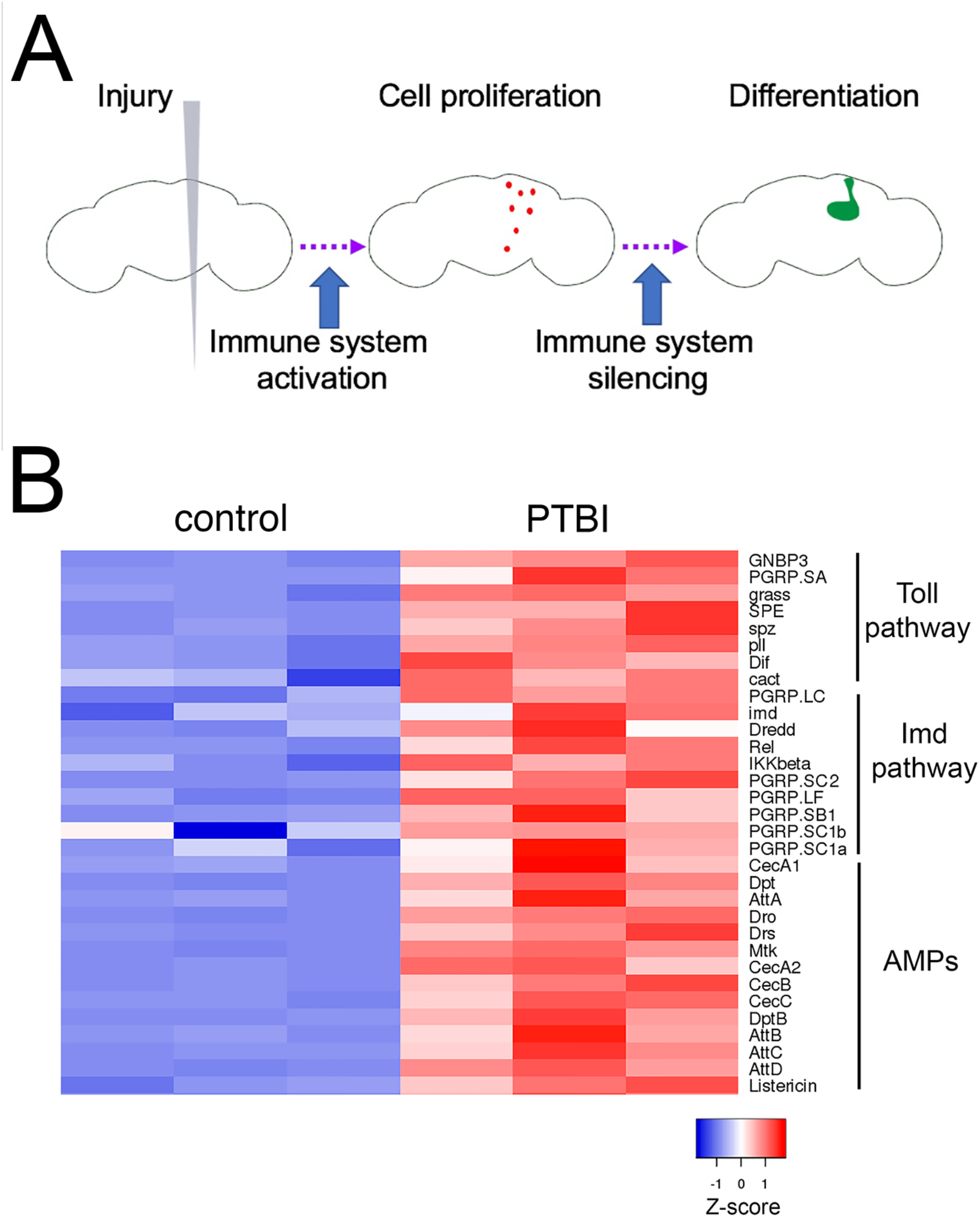
Penetrating Traumatic Brain Injury (PTBI) induces cell proliferation and adult neurogenesis via activation of the immune system. **A**. The immune system is activated acutely and transiently after PTBI. We propose that this transient activation triggers proliferation of cells (red) particularly around the area of injury. The immune system is then silenced, as evidenced by the downregulation of innate immune pathway genes. The newly created cells can differentiate into new neurons and glia that go on to integrate and repair the damage to the mushroom body (green). **B**. Heat map comparing expression of innate immunity genes from 4-hour post-injury RNA-Seq data. The three control replicates are on the left, and the injured samples are on the right. Red indicates upregulation, while blue indicates downregulation. Multiple components of the Toll and Imd signaling pathways were upregulated. The canonical downstream targets of Toll and Imd signaling, the antimicrobial peptides (AMPs) also are upregulated by PTBI. This heat map plots Z-scores, which represent the number of standard deviations the expression for each sample is from the mean expression of all samples analyzed.

### Mutants in NF-κB Transcription Factor Genes Display Proliferation Defects After Injury

To test whether Toll or Imd signaling is necessary after injury for cell proliferation, we utilized mutants for the major effectors of signaling in each pathway. As both pathways converge on NF-κB transcription factors, we used allelic combinations of mutants for *Dif* and *Rel* to block Toll and Imd signaling, respectively. We injured the mutant flies using our standard PTBI technique and allowed the flies to recover for 24 hours before assaying for cell proliferation. We used two methods to detect cell proliferation. The first was using an antibody to stain for phospho-histone H3 (PH3), a histone variant present during the final stages of S phase through telophase of M phase (Hans and Dimitrov, 2001). We found that in animals of our control genotype there was the stereotypical increase in the number of proliferating cells seen in injured brains as compared to uninjured controls (**Fig. 2A-BB, I**). However, in injured *Dif* or *Rel* mutants, there was not an increase in the number of PH3^+^ cells after injury (**Fig. 2C-FF,I**). We also assayed flies that were mutant for both *Dif* and *Rel*, and we observed a similar lack of injury-stimulated proliferation these samples (**Fig. 2G-HH, I**). Because PH3 is only present for a limited portion of the cell cycle, we were concerned that we these assays would not capture the scope of proliferation. We therefore repeated these experiments using a second cell proliferation labeling technique, 5-ethynyl-2’-deoxyuridine (EdU), which is incorporated into newly synthesized DNA to permanently label newly created cells and their daughters. We observed the same trend using this labeling technique, with fewer proliferating cells in the *Dif* and *Rel* mutants after injury as compared to the injured control brains. We likewise saw no significant difference between control and injured *Dif; Rel* double mutants, indicating that the activation of proliferation in response to injury does not occur in the double mutant background (**Fig. S1**). We conclude that both Toll and Imd signaling are required for cell proliferation after PTBI. Because the numbers of proliferating cells detected were similar with anti-PH3 and EdU labeling, we chose to use PH3 staining to detect proliferating cells for the remainder of the experiments described here.

**Figure 2.**
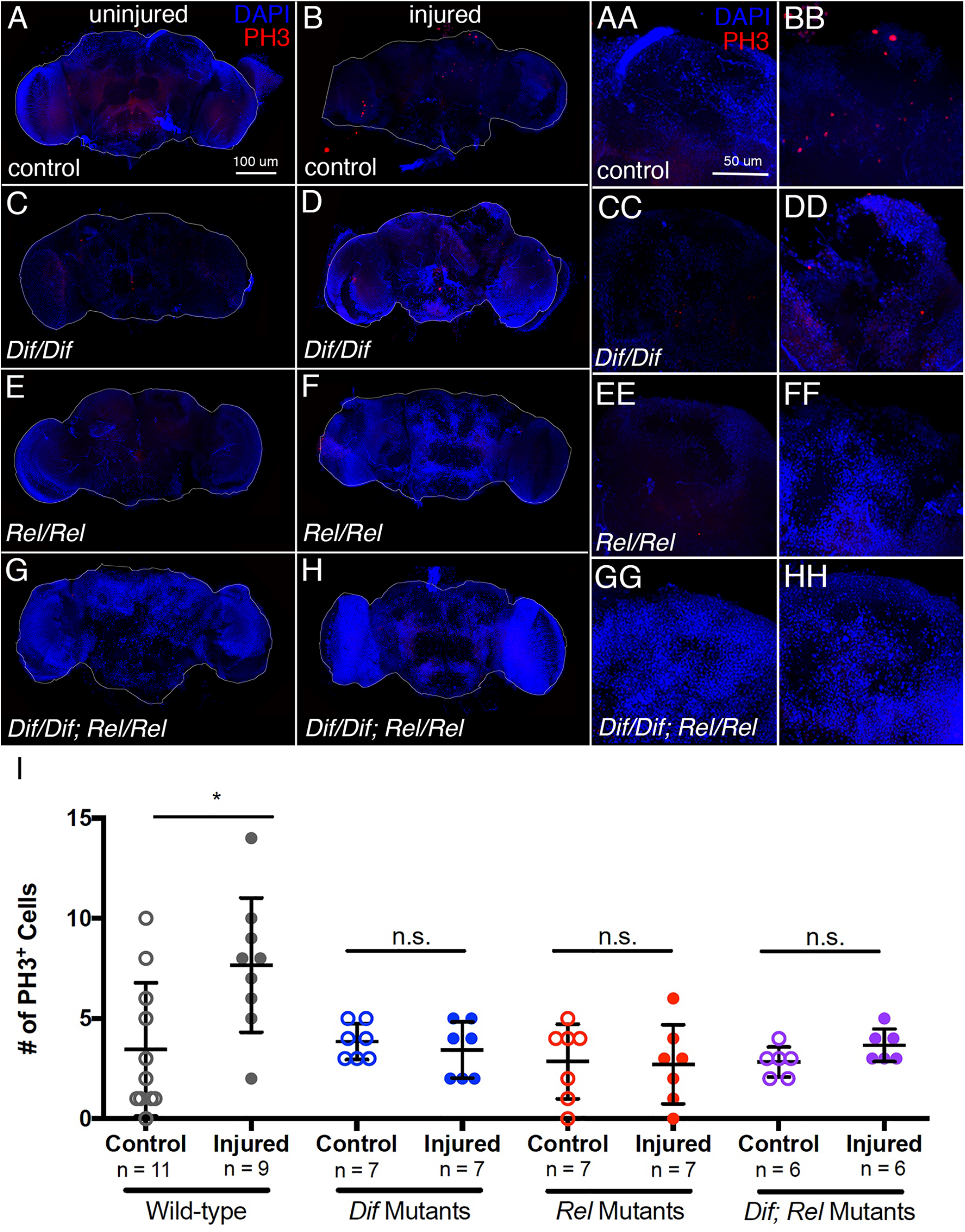
*Dif* and *Rel* mutants do not exhibit increased cell proliferation following PTBI. **A-H**. Cell proliferation assayed by anti-PH3 labeling (in red) 24 hours after injury reveals that while baseline cell proliferation (**A**) is not significantly reduced in uninjured *Dif* (**C**), *Rel* (**E**) and *Dif;Rel* (**G**) mutants, cell proliferation is not stimulated by PTBI in the mutants (**D**,**F**,**H**) as it is in control (**B**) animals. The right mushroom bodies of each of these brains are shown in higher magnification in **AA-HH. I**. Quantification of cell proliferation in *Dif* and *Rel* mutants reveals no statistically significant increase after injury. Wild-type control brains had an average of 3.5 PH3+ cells per brain, and wild-type injured samples had 7.7 (p-value = 0.0117). *Dif* mutant control brains had 3.9 PH3+ cells per brain, and injured samples had 3.4 PH3+ cells (p-value = 0.5080). *Rel* mutant control brains had 2.9 PH3+ cells on average, and injured brains had 2.7 PH3-positive cells (p-value = 0.8917). The *Dif; Rel* double mutant control samples had 2.8 PH3-positive cells per brain on average, and the injured samples had 3.7 PH3-positive cells (p-value = 0.0959). Error bars reflect SD.

### Immune system activation is transient after PTBI

Previous work described in Cao *et al*., 2013 showed that chronic activation of the immune system and expression of AMPs in the brain resulted in neurodegeneration (Cao et al., 2013). We therefore wanted to test whether the levels of the AMPs were chronically activated post-PTBI. We conducted qRT-PCR to measure levels of four AMPs at 2, 4, 12, and 24 hours after injury (**Fig. 3A**). Expression levels of all four AMPs peaked at 12 hours after injury, with 20 times more *CecB*, 228 times more *DptA*, 97 times more *Dro*, and 2.6 times more *Drs* expression in injured samples than controls. The levels of transcripts for all of these genes returned toward baseline 24 hours after injury, indicating that the peak of expression is early after PTBI. Taken together these data show that the immune system is rapidly and acutely activated upon injury but is quickly suppressed. This temporal dynamic may play an important role in inducing proliferation and regeneration as opposed to the degeneration seen in other studies where there is chronic expression of these genes.

**Figure 3.**
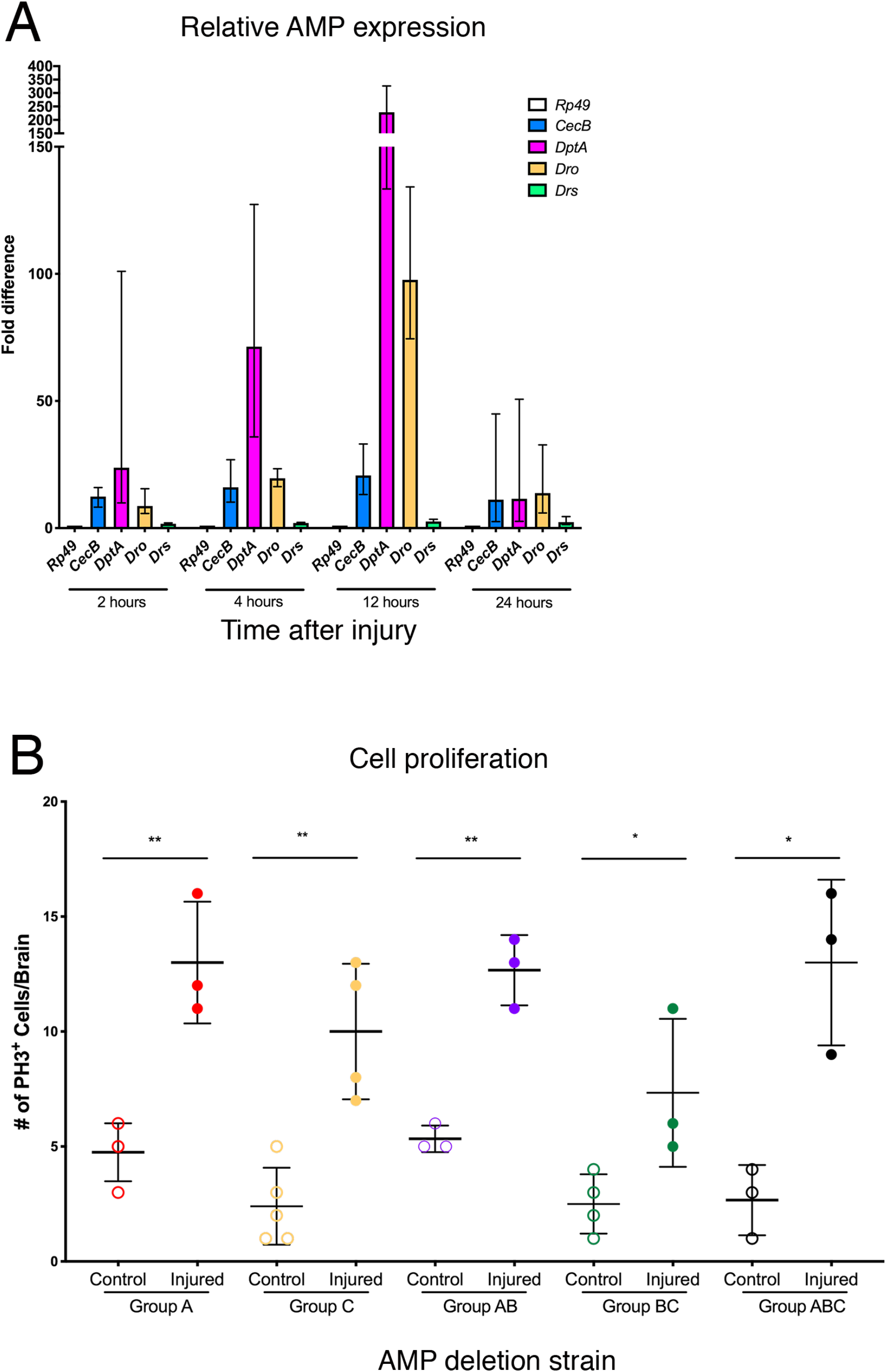
Antimicrobial peptides (AMPs) are not required for the proliferative response. **A**. AMP gene expression is upregulated following PTBI. The mRNA levels of 4 different AMPs were assayed at 2, 4, 12 and 24 hours after injury. The level of *CecB* mRNA is increased more than 10-fold by 2 hours post-PTBI and remains elevated at 24 hours. In contrast, the levels of *DptA* and *Dro* mRNAs increase 100-200-fold with a peak at 12 hours and a return to near baseline by 24 hours post-PTBI. The level of *Drs* mRNA increases only ∼2-fold by 2 hours post-PTBI and remains slightly elevated at 24 hours. The qRT-PCR results reflect triplicate biological samples, represented relative to the levels of *Rp49*, and then normalized to the corresponding levels in time-matched controls. Error bars calculated by Relative Expression Software Tool analysis and reflect SEM. Note that scales on Y axes differ among the graphs. **B**. Fly stocks lacking groups of AMP genes and combinations of these groups (see text for details) were assayed for cell proliferation 24 hours post-injury using PH3 staining. Loss of any single group of AMPs was not sufficient to reduce cell proliferation after injury. Nor did pairwise combinations of AMP deletions (Group AB, Group BC) exhibit defects in cell proliferation post-PTBI. Finally, flies carrying all three groups of AMP deletions had no defects in cell proliferation. Error bars represent SD. Group A control brains had 4.8 PH3-positive cells per brain, and injured samples had 13 PH3-positive cells (p-value = 0.0026). Group C control samples had 2.4 PH3-positive cells, and injured samples had 10 PH3-positive cells (p-value = 0.0017). Group AB control flies had 5.3 PH3-positive cells, and injured samples of the same genotype had 12.7 PH3-positive cells per brain (p-value = 0.0015). Group BC control brains had 2.5 PH3-positive cells, and injured brains had 7.3 PH3-positive cells (p-value = 0.0383). Group ABC control brains had 2.7 PH3-positive cells per brain while injured samples had 13.00 PH3-positive cells (p-value = 0.0103). At least 3 brains were imaged for each condition.

### Canonical Targets of Toll and Imd Signaling Are Not Cell Proliferation Effectors

AMPs are a standard readout of innate immune system activation in *Drosophila* and can coordinate cellular responses, including neurodegeneration (Cao et al., 2013). If AMPs are required for cell proliferation after injury, we would expect to see a reduction in the number of proliferating cells when AMP expression cannot be stimulated. Hanson *et al*. have grouped the 14 AMPs into three groups and generated compound mutants for 10 of these (Cao et al., 2013). Group A is composed of *Defensin* (*Def*) and the *Cecropins* (*Cec*) and is primarily regulated by the Imd pathway, Group B contains *Drosocin* (*Dro*), the *Diptericins* (*Dipt*) and the *Attacins* (*Att*) and are regulated primarily by the Imd pathway, and Group C has *Metchnikowin* (*Mtk*) and *Drosomycin* (*Drs*) and is controlled by the Toll pathway (Hanson et al., 2019). We subjected Groups A, C, AB, BC and ABC flies to our standard PTBI protocol and assayed for cell proliferation using the anti-PH3 antibody 24 hours after injury. We found that no combination of deletion groups, including loss of all three, reduced the increase in PH3-positive cells seen after injury (**Fig. 3B**). These results indicate that these 10 AMPs are not the targets of Toll or Imd signaling required for triggering cell proliferation post-PTBI and that there may be non-canonical targets that play an essential role in stimulating regeneration in adult brains. Because these compound mutants did not include the *Cecropin* genes, we cannot rule out a function for Cecropins in activating cell proliferation post-PTBI. Nonetheless, this seems unlikely given the upregulation of *CecB* expression post-PTBI in both *Dif* and *Rel* mutants (**Fig. S2**), both of which lack a proliferative response after injury (**Fig. 2**). Also, although the baseline expression of multiple AMPs is reduced in *Dif* and *Rel* mutants, AMP expression is nonetheless stimulated by injury (**Fig. S2**). This is consistent with the hypothesis that non-canonical targets of Dif and Rel activate cell proliferation after PTBI.

### Toll and Imd Signaling Are Required in Specific Tissues for Cell Proliferation after Injury

There are multiple tissues present in the *Drosophila* head capsule that are affected by PTBI. Specifically, in addition to neurons and glia in the brain, two tissues known to be important to the innate immune response are located in the head. These are the fat body, a major site of AMP production following injury and infection (Aggarwal and Silverman, 2008; Kounatidis and Ligoxygakis, 2012; Lemaitre and Hoffmann, 2007), and the hemocytes, which phagocytose debris, and also produce AMPs (Williams, 2007). To determine the requirements for Toll and Imd signaling in particular tissues, we designed tissue-specific knockdown experiments utilizing the GAL4-UAS binary system (Brand and Perrimon, 1993) to specifically reduce *Dif* or *Rel* expression via RNA interference (RNAi). We reduced Dif and Rel levels throughout the animal (*Actin-GAL4*), in neurons (*C155-GAL4*), in glia (*Repo-GAL4*), in fat body (*yolk-GAL4*), or in hemocytes (*Hml-GAL4*) and compared the number of proliferating cells per brain after PTBI to controls (**Fig. 4A, B**, quantification in **Table S1** and **Table S2**). The ubiquitous knockdown of either *Dif* or *Rel* is sufficient to reduce the number of proliferating cells after injury compared to injured controls (**Fig. 4A, B**), phenocopying the results we obtained from the mutants (**Fig. 2**). We found that knockdown of neither *Dif* nor *Rel* in neurons significantly reduced the amount of cell proliferation after injury (**Fig. 4B**), suggesting that immune signaling is not required in neurons for proliferation after injury. In contrast, immune signaling via both Toll and Imd pathways is required in both glia and fat body, as there was a significant reduction in the number of PH3+ cells when either *Dif* or *Rel* were knocked down in either tissue (**Fig. 4B**). Interestingly, *Rel* knockdown in hemocytes was sufficient to prevent cell proliferation after injury (**Fig. 4B**), suggesting that Imd and Toll may be involved in different processes in distinct tissues. Together, these data indicate that the innate immune pathways are required in multiple tissues both within the damaged brain and outside of it to coordinate the activation of cell proliferation that will generate new neural tissue.

**Figure 4.**
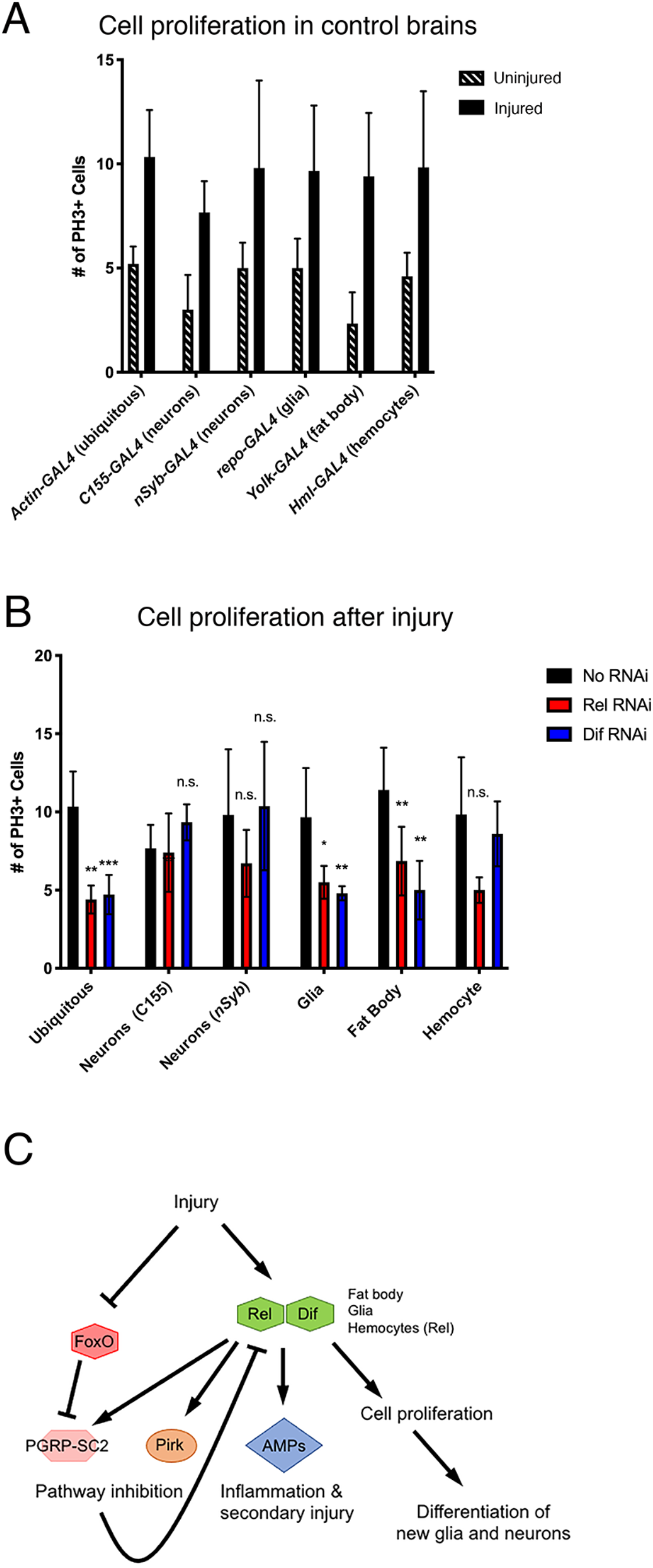
Tissue specific knockdown of *Dif* and *Rel* reveals different requirements for the Toll and Imd pathways in activating cell proliferation following PTBI. **A, B**. To test where the Toll and Imd innate immunity pathways are required, *Dif* or *Rel* expression were knocked down using UAS-RNAi transgenes. Ubiquitous knockdown was achieved with *Actin-GAL4*, neuronal knockdown with *nSyb-GAL4* and *C155-GAL4*, glial knockdown with *repo-GAL4*, fat body knockdown with *Yolk-GAL4* and hemocyte knockdown with *Hml-GAL4*. Controls for this experiment were Drosophila harboring the GAL4 drivers but lacking RNAi constructs (**A**). There was a significant increase in the number of proliferating cells after injury in all of the control strains (**A**). However, there was reduced cell proliferation after injury when *Dif* expression was reduced ubiquitously, in glia, or in the fat body (**B**). There also was reduced cell proliferation after injury when *Rel* expression was reduced ubiquitously, in glia, or in the fat body (**B**). However, Rel knockdown in hemocytes also showed an effect (**B**). Error bars reflect SD. At least five brains were imaged per condition and genotype, and the numbers of PH3-positive cells are included in **Tables S1** and **S2. C**. Proposed model for the regulation of cell proliferation after PTBI. Injury stimulates the Toll and Imd pathways, resulting in increased expression of *Dif* and *Rel*. Injury leads to downregulation of *FoxO* via an unknown mechanism. Reduction of FoxO leads to derepression of the Imd pathway inhibitor PGRP-SC2 (Guo et al., 2014). Dif and Rel activate the AMP genes and the pathway repressors *pirk* and *PGRP-SC2*. Pirk and PGRP-SC2 negatively regulate the Imd pathway, leading to downregulation of the *AMP* genes by 24 hours post-PTBI. *AMP* downregulation is essential to prevent cell death and neurodegeneration. Injury also stimulates cell proliferation via non-canonical Dif and Rel targets, leading to the generation of new glia and new neurons. We note that FlyBase recently published new guidelines for the standardization of signaling pathway components (http://flybase.org/lists/FBgg/pathways). These conventions are followed in the model.

## Discussion

Neural regeneration after injury is a complex process that likely involves multiple pathways to coordinate cell proliferation, differentiation, and integration of cells into proper circuits. To begin to dissect the mechanisms underlying neural regeneration, we focused on the initial cell proliferation response that occurs within 24 hours of injury. We found that there is a rapid and dramatic induction of both the Toll and Imd innate immunity pathways prior to cell division. Toll and Imd have been related to repair responses in the embryonic epidermis and larval tumor systems (Capilla et al., 2017; Parisi et al., 2014), and we wanted to test whether these pathways were similarly required for neural injury response. Mutants for either of the NF-κB effectors of the Toll and Imd pathways, *Dif* and *Rel* respectively, display defects in the induction of cell proliferation after injury, indicating that both Toll and Imd signaling are indeed necessary for the initial injury response. Signaling is required specifically in the fat body and glia for both pathways, but there is an additional role for Imd activation in hemocytes, indicating a requirement for signals from both within the injured brain and outside of it. We also found that many of the AMPs, the major readouts of immune signaling, are dispensable for activating cell proliferation. We therefore propose that non-canonical targets are responsible for this aspect of Toll and Imd signaling.

There is precedence for innate immune signaling in regulating tissue repair after injury. For instance, injury of the *Drosophila* embryonic epidermis activates Toll signaling, which is required to induce the barrier repair genes (Carvalho et al., 2014). In *Drosophila* larvae, Toll signaling is triggered by hemocytes that have been recruited to tumors or damaged regions and coordinates with other pathways involved survival and death, such as the c-Jun N-terminal kinase (JNK) pathway (Parisi et al., 2014). Additionally, the Tak1/Tab complex that regulates phosphorylation of Rel has been shown to regulate JNK signaling by activating Hemipterous (Hep). Hep is the homolog of MKK7, which phosphorylates basket (Bsk), the *Drosophila* homolog of JNK (Parisi et al., 2014). In our 4 hour post-PTBI RNA-Seq data, we observed increased expression of JNK components. We also observed upregulation of activators of the Janus kinase/signal transducers and activators of transcription (JAK/STAT) pathway, particularly Unpaired 2 (Upd2) and Unpaired 3 (Upd3). A major downstream effector of JAK/STAT signaling is the transcription factor STAT92E. For one pro-survival target, *Turandot A*, both Rel and STAT92E are required for gene expression (Ekengren and Hultmark, 2001).

Thus, it is reasonable to hypothesize that Toll and Imd are working in coordination with JNK and/or JAK/STAT pathways to stimulate a regenerative response in the injured brain. Further, the activation of distinct signaling pathways may depend on activity both in the damaged brain and in other tissues, including the fat body. It therefore will be important to investigate the extent to which these other pathways contribute to the regenerative response, by first determining if they are required for proliferation to occur after PTBI and subsequently testing whether they are needed in the same or different tissues than Toll and Imd. This could provide insight into why therapies that rely on activation of only one pathway are not sufficient to induce the repair of damaged tissue.

One question that arises from our findings is what is the role of AMPs if not to stimulate cell proliferation? Cao *et al*., 2013 reported AMP overexpression was sufficient to induce neurodegeneration through unknown mechanisms (Cao et al., 2013), leading us to hypothesize that AMPs may play a similar role to toxic cytokines or neuroinflammation in mammalian systems to promote secondary injury and cell death. Studies utilizing a closed head model of TBI in *Drosophila* reported a sustained increase in the expression of innate immunity genes, and these studies also report substantial mortality after injury (Katzenberger et al., 2013). In contrast, the expression of AMPs in the PTBI model is transient, and the mortality is negligible at equivalent times after injury (Crocker et al., 2021). Together, these data support the idea that while prolonged expression of AMPs is deleterious, transient expression may be beneficial.

A critical question raised by this work is how transient activation of the Toll and Imd pathways is achieved. Our RNA-Seq data offer a clue. Specifically, the expression of two negative regulators of the Imd pathway is rapidly and significantly elevated after PTBI. These negative regulators are Pirk and PGRP-SC2 (Kleino et al., 2008; Paredes et al., 2011). Pirk acts intracellularly to downregulate Rel activity. PGRP-SC2 is secreted and therefore can downregulate the Imd pathway both cell autonomously and non-cell autonomously, offering the possibility of direct crosstalk among tissues. We hypothesize that these negative regulators play an essential role in the rapid downregulation of the innate immune system following PTBI, thereby preventing cell death and neurodegeneration (**Fig. 4C**) and facilitating neurogenesis. Both Pirk and PGRP-SC2 are known targets of Rel. Thus, activation of the Imd pathway can be self-limiting. However, neither Pirk nor PGRP-SC2 are upregulated following a closed head TBI (Katzenberger et al., 2016). One possible explanation for this is the upregulation of FoxO. FoxO is a known repressor of PGRP-SC2 (Guo et al., 2014) and is slightly downregulated in our four hour post-PTBI RNA-Seq dataset. We propose that the reduction in FoxO levels combines with an increase in Rel activity to activate expression of Imd pathway inhibitors, including PGRP-SC2 (**Fig. 4C**).

It was long thought that immune activation and inflammation after CNS injuries or onset of neurodegenerative disorders had exclusively negative consequences. Activation of glia is associated with high levels of cytokine production and blood-brain barrier (BBB) permeability, which compromises animal health (DiSabato et al., 2016). In particular, neuroinflammation was reported to reduce the proliferation of NSCs and decrease survival and integration of newly created neurons (Ekdahl et al., 2003). However, more recent experiments have uncovered a role for innate immune signaling in promoting repair. Mice that lack IL-6 or TNF, immune signals that are associated with the inflammatory response, have a higher mortality after closed head injury, indicating that there may be a protective role for immune system activation after TBI (Morganti-Kossmann et al., 2002). After damage to the olfactory neurons, TNF-α and its downstream effector RelA, a mammalian homolog of the *Drosophila* Relish protein, are required in the injured cells to trigger the proliferation of nearby cells to repair damage (Chen et al., 2017). Consistent with this, transient innate immune activation also has been associated with neurogenesis in mammalian systems (Morganti-Kossman et al., 1997).

Our *Drosophila* PTBI model may help to uncover links between the immune system and neural regeneration that also are present in mammals. This work is a start to dissecting how the same innate immune pathways can contribute to both regenerative and degenerative phenotypes following neural injury. Further analysis of the mechanisms underlying modulation of the innate immune response following PTBI are like to reveal how we might manipulate these pathways to shift the balance from negative effects to more beneficial ones (Morganti-Kossman et al., 1997). This work demonstrates the first direct links between the induction of neural regeneration and innate immune signaling in *Drosophila* and supports evidence linking immune signaling in mammalian brains to proliferation and regeneration.

## Acknowledgements

We are grateful to Becky Katzenberger and Sarah Neuman for technical assistance. We also would like to thank Barry Ganetzky and David Wassarman for lively discussions that undoubtedly improved the science and Kent Mok, Cayla Guerra, and Bailey Spiegelberg for their contributions to the laboratory. Most of the *Drosophila* strains used in this study were obtained from the Bloomington Drosophila Stock Center (BDSC; NIH P40OD018537). This work was supported by NIH T32 GM007133 (KM and KLC); NIH NS090190 (GBF); NIH NS102698 (GBF); the University of Wisconsin Graduate School (GBF); and the Women in Science and Engineering Leadership Institute (WISELI) (GBF).

## Figure Legends

**Figure S1.**
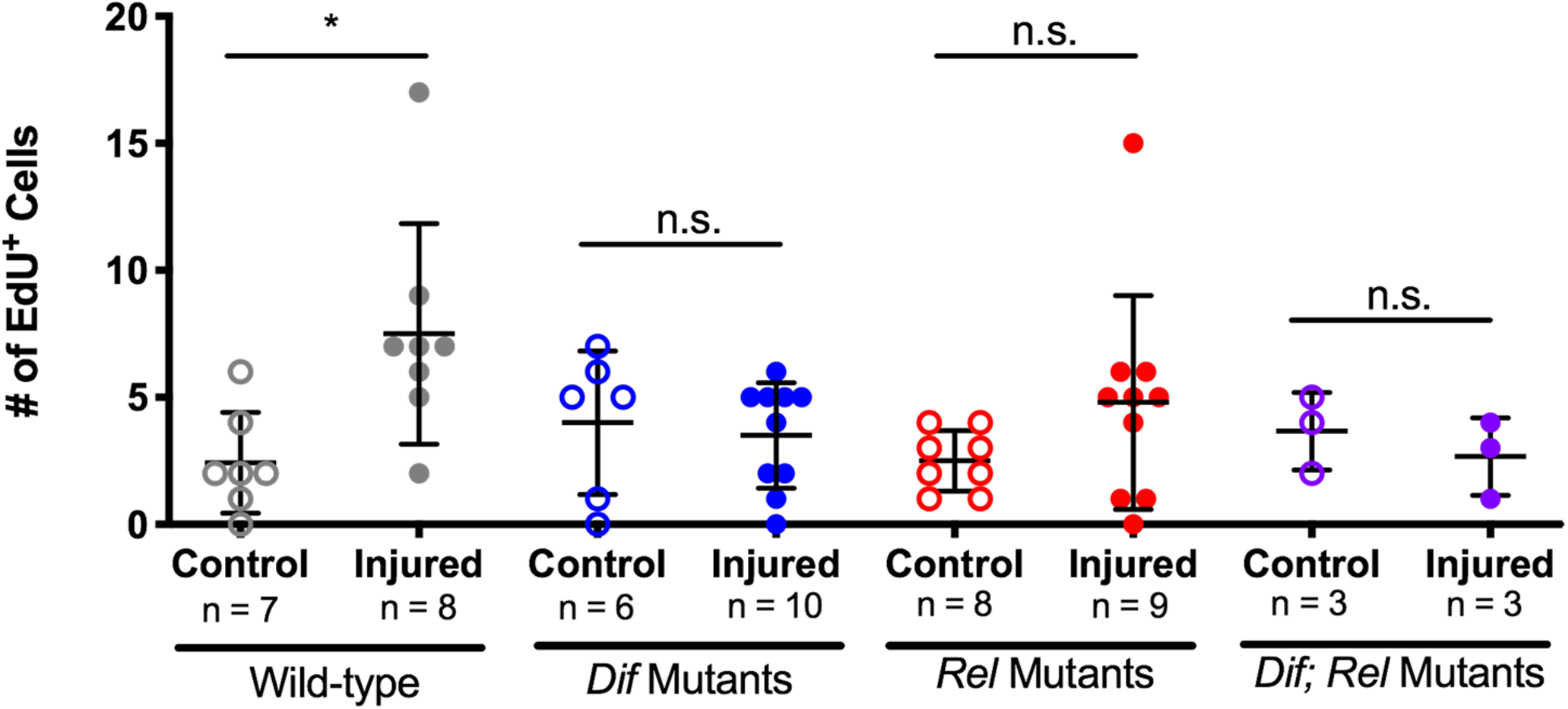
*Dif* and *Rel* mutants do not exhibit increased cell proliferation following PTBI when assayed with EdU 24 hours after injury. Cell proliferation assayed by EdU incorporation reveals similar results to anti-PH3 staining (**Fig. 2**). While baseline cell proliferation is not significantly reduced in uninjured *Dif, Rel* and *Dif; Rel* mutants, cell proliferation is not stimulated by PTBI in the mutants as it is in control animals. Error bars reflect SD. Wild-type control brains had 2.4 EdU-positive cells per brain, and the injured samples had 7.1 EdU-positive cells (p-value = 0.0115). *Dif* mutant control brains had 4 EdU-positive cells per brain on average, and *Dif* mutant injured brains had 3.5 EdU-positive cells on average (p-value = 0.6889). The *Rel* mutant controls had 2.5 EdU-positive cells per brain, and the injured ones had 3.7 EdU-positive cells (p-value = 0.225). The *Dif; Rel* double mutant controls had 3.7 EdU-positive cells on average, and the injured samples had 2.7 EdU-positive cells (p-value = 0.4676).

**Figure S2:**
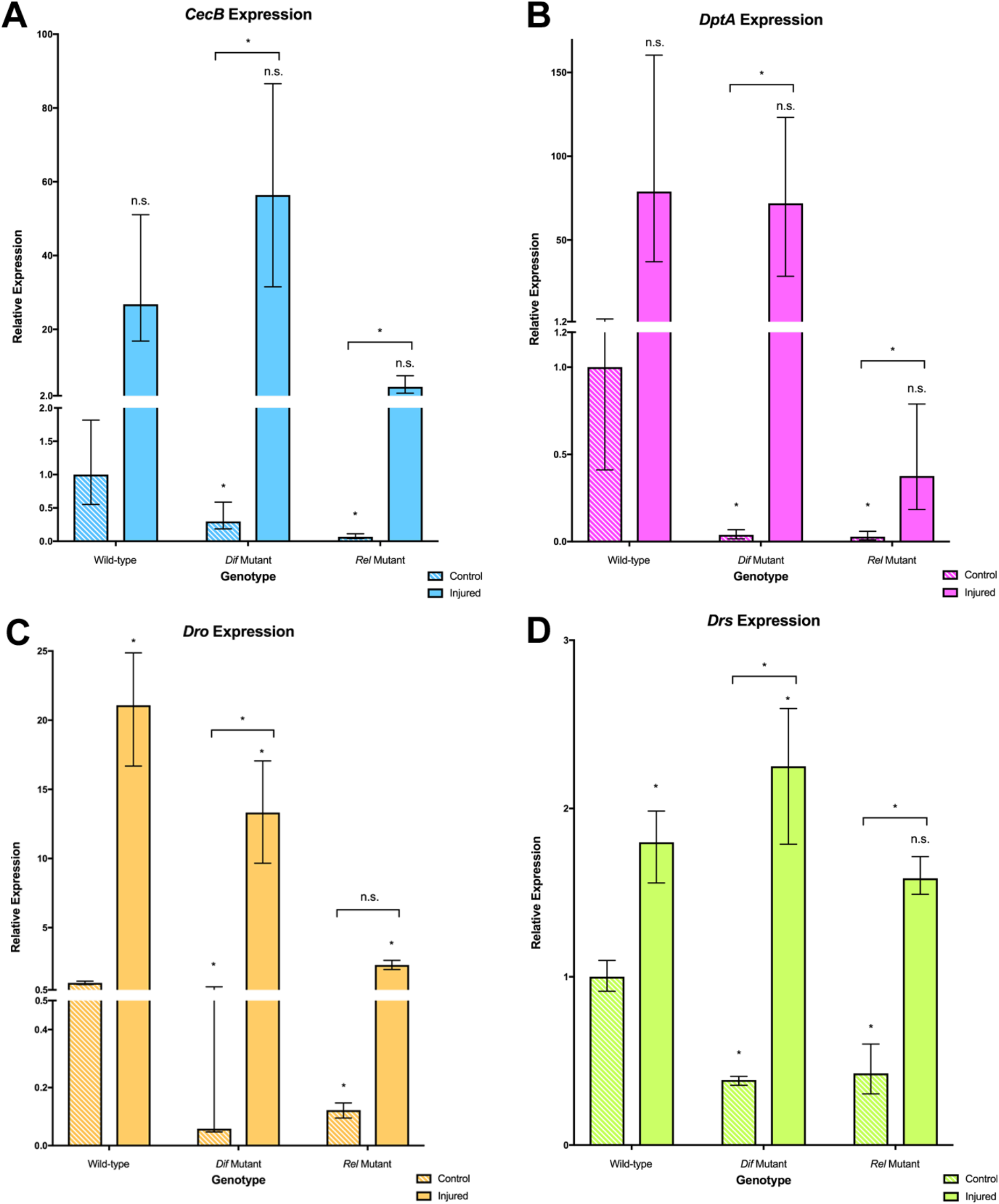
Expression of AMPs is reduced in both uninjured and injured *Dif* and *Rel* mutant samples. The mRNA levels of 4 different AMPs were assayed in six conditions: control and injured wild-type (*OK107 > GFP*), control and injured *Dif* mutants, and control and injured *Rel* mutants. All RNA samples were collected four hours after injury. We found that baseline expression of all AMPs was significantly reduced in the control *Dif* and *Rel* mutant samples, indicating that *Dif* and *Rel* play essential roles in maintaining baseline expression of these genes. After injury, we see an increase in expression of all four AMPs in all conditions. Loss of *Dif* does not dramatically lower expression of these genes after injury, as relative expression of *CecB, DptA*, and *Drs* are all nearly equal to or greater than the relative expression seen in wild-type injured samples. Loss of *Rel* does seem to reduce the relative expression of all four genes after injury, indicating that the Imd pathway may play a more integral role in regulating AMP expression. The qPCR results reflect triplicate biological samples, represented relative to the levels of *Rp49*, and then normalized to the corresponding levels in the 4hr wild-type control samples. Error bars calculated by Relative Expression Software Tool analysis and reflect SEM. Note that scales on Y axes differ among the graphs.

**Table S1.** Quantification of PH3-positive cells per brain in knockdown experiments.

| **Genotype** | **Average of PH3-positive cells per control brain** | **Average of PH3-positive cells per injured brain** | **Significance** |
| --- | --- | --- | --- |
| *Actin-GAL4* | 5.2 | 10.3 | Yes (p-value = 0.0099) |
| *Actin>Dif RNAi* | 3.4 | 4.7 | No |
| *Actin>Rel RNAi* | 4.0 | 4.4 | No |
| *Hml-GAL4* | 4.6 | 9.8 | Yes (p-value = 0.0137) |
| *Hml>Dif RNAi* | 3.6 | 8.6 | Yes (p-value = 0.0015) |
| *Hml>Rel RNAi* | 4.3 | 5.0 | No |
| *nSyb-GAL4* | 5.0 | 9.8 | Yes (p-value = 0.039) |
| *nSyb>Dif RNAi* | 3.5 | 10.4 | Yes (p-value = 0.0019) |
| *nSyb>Rel RNAi* | 2.6 | 6.7 | Yes (p-value = 0.0030) |
| *Repo-GAL4* | 5.0 | 9.7 | Yes (p-value = 0.0078) |
| *Repo>Dif RNAi* | 2.6 | 4.8 | Yes (p-value = 0.0039) |
| *Repo>Rel RNAi* | 3.4 | 5.5 | Yes (p-value = 0.0395) |
| *Yolk-GAL4* | 2.3 | 9.4 | Yes (p-value = 0.0007) |
| *Yolk>Dif RNAi* | 3.0 | 5.0 | No |
| *Yolk>Rel RNAi* | 3.7 | 6.9 | Yes (p-value = 0.0016) |
| *C155-GAL4* | 3.0 | 7.7 | Yes (p-value = 0.0005) |
| *C155>Dif RNAi* | 3.8 | 9.3 | Yes (p-value = 0.0008) |
| *C155>Rel RNAi* | 3.3 | 7.4 | Yes (p-value = 0.0196) |

**Table S2.**
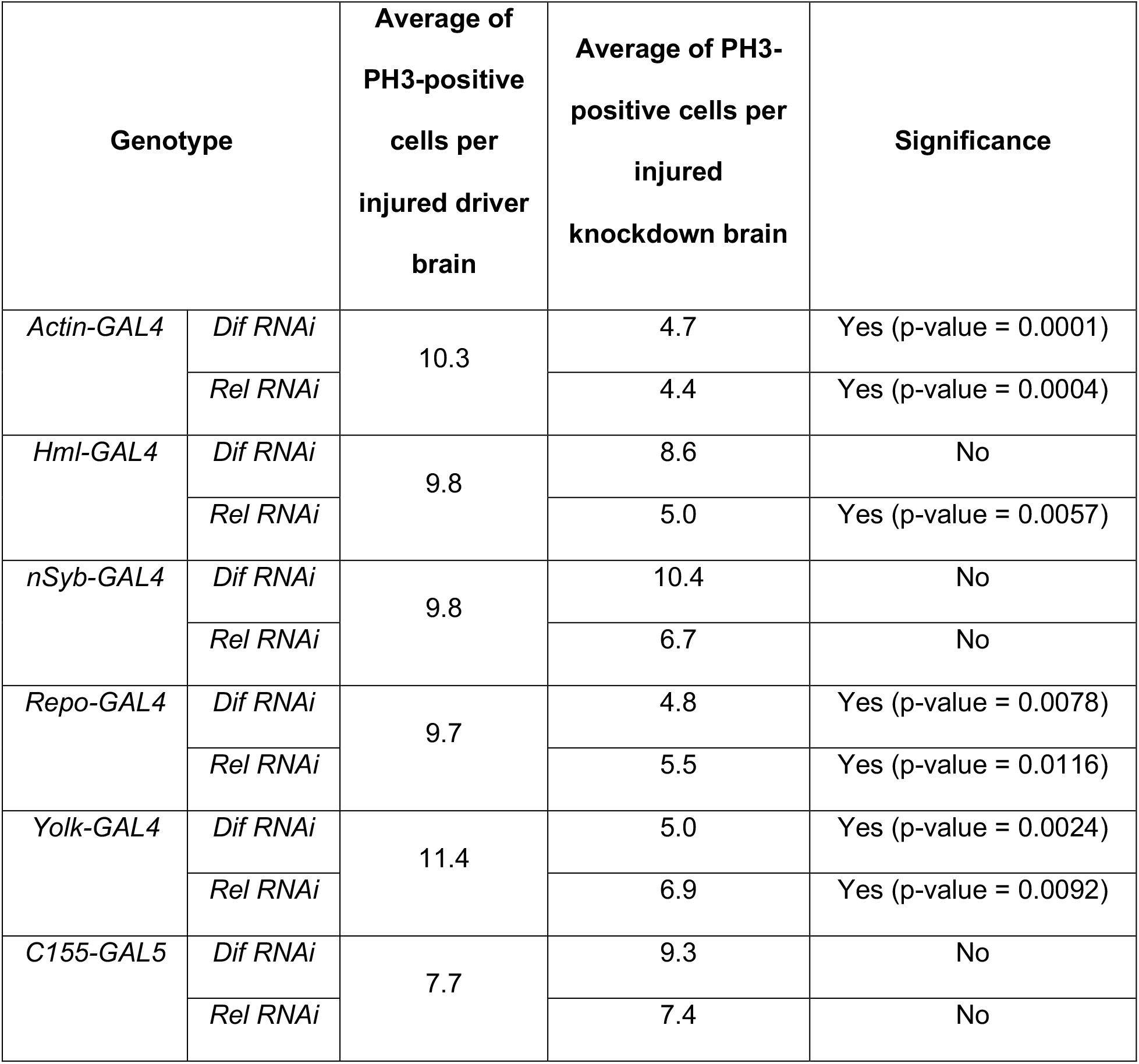
Quantification and Comparison PH3-positive Cells in Injured *GAL4* Driver and Injured RNAi Knockdown Samples.

